# A dispensable role of mitochondrial fission protein 1 (Fis1) in the erythrocytic development of *Plasmodium falciparum*

**DOI:** 10.1101/2020.06.18.160663

**Authors:** Mulaka Maruthi, Liqin Ling, Jing zhou, Hangjun Ke

**Author notes:** Address correspondence to Dr. Hangjun Ke. Department of Biochemistry, Central University of Haryana, Haryana, India-123031 (Mulaka Maruthi).

## Abstract

Malaria remains a huge global health burden and control of this disease has run into a severe bottleneck. To defeat malaria and reach the goal of eradication, a deep understanding of parasite biology is urgently needed. The mitochondrion of the malaria parasite is essential throughout the parasite’s lifecycle and has been validated as a clinical drug target. In the asexual development of *Plasmodium spp*., the single mitochondrion grows from a small tubular structure to a complex branched network. At the end of schizogony when 8-32 merozoites are produced, the branched mitochondrion is precisely divided, distributing one mitochondrion to each forming daughter merozoite. In mosquito and liver stages, the giant mitochondrial network is split into thousands of pieces then daughter mitochondria are segregated into individual progeny. Despite the significance of mitochondrial fission in *Plasmodium*, the underlying mechanism is largely unknown. Studies of mitochondrial fission in model eukaryotes have revealed that several mitochondrial fission adaptor proteins are involved in recruiting dynamin GTPases to physically split mitochondrial membranes. Apicomplexan parasites, however, share no identifiable homologs of mitochondrial fission adaptor proteins of yeast or human, except for Fis1. Here, we investigated the localization and essentiality of the Fis1 homolog in *Plasmodium falciparum*, PfFis1 (PF3D7_1325600), during the asexual lifecycle. We found that PfFis1 requires an intact C-terminus for mitochondrial localization but is not essential for parasite development or mitochondrial fission. The dispensable role of PfFis1 indicates *Plasmodium* contains additional fission adaptor proteins on the mitochondrial outer membrane that could be essential for mitochondrial fission.

**Importance:** Malaria is responsible for over 230 million clinical cases and ∼ half a million deaths each year. The single mitochondrion of the malaria parasite functions as a metabolic hub throughout the parasite’s developmental cycle as well as a source of ATP in certain stages. To pass on its essential functions, the parasite’s mitochondrion needs to be properly divided and segregated into all progeny during cell division via a process named mitochondrial fission. Due to the divergent nature of *Plasmodium spp*., molecular players involved in mitochondrial fission and their mechanisms of action remain largely unknown. We found that Fis1, the only identifiable mitochondrial fission adaptor protein evolutionarily conserved in the phylum of Apicomplexa, however, is not essential for *Plasmodium falciparum*. Our data suggest that malaria parasites use redundant fission adaptor proteins on the mitochondrial outer membrane to mediate the fission process.

---

Malaria is a major cause of human morbidity and mortality globally (1). The clinical symptoms of malaria are mainly resulted from repetitive growth of *Plasmodium* parasites inside the red blood cells (RBCs) and continuous rupture of infected RBCs. Post invasion of a host cell, the parasite lives inside the parasitophorous vacuole, undergoing growth and division to generate new infective progeny. Unlike many cells that divide via binary fission, inside RBC, the malaria parasite replicates its nuclear DNA 3-5 rounds without concurrent cytokinesis, resulting in the formation of 8-32 daughter cells (merozoites). Division of 8-32 merozoites starts at the very end of the lifecycle. This unique reproduction manner of *Plasmodium* during the asexual blood stage is termed schizogony (2). In mosquito and liver stages, the parasite’s nuclear DNA is replicated 13-14 rounds without cytokinesis, producing up to 10,000 progenies all at once (3); this process is termed sporogony. To coordinate this unique cellular reproduction mechanism (schizogony or sporogony), the single mitochondrion of the malaria parasite also undergoes a process of growth and division. In the asexual blood stage of *Plasmodium falciparum*, the mitochondrion grows from a single small tubular structure to a large branched network over the 48 h intraerythrocytic development cycle (IDC) (4), which is then divided into 8-32 pieces to provide one merozoite with one daughter mitochondrion. Since the mitochondrion is essential for parasite growth and replication and cannot be made *de novo*, the process of mitochondrial fission is critical to the malaria parasite.

Studies on mitochondrial fission in model organisms have suggested that, the fission machinery that “splits” mitochondrial membranes involves at least two classes of molecules, mitochondrial fission adaptor proteins located on the mitochondrial outer membrane (MOM) and fission GTPases (mechanoenzymes) that are recruited from cytosol to the mitochondrial fission sites (5). In mammalian cells, multiple mitochondrial fission adaptor proteins have been identified to recruit the fission GTPase (dynamin related protein 1, Drp1), including Fis1 (mitochondrial fission protein 1), Mff (mitochondrial fission factor), MiD49 and MiD51 (mitochondrial dynamics proteins 49 kDa and 51 kDa) (6). Interestingly, except for Fis1, other mammalian MOM-bound fission adaptor proteins (Mff, MiD49 and MiD51) have not been identified in yeast (*Saccharomyces cerevisiae*) (7), plants (7), or unicellular protozoans, suggesting that mitochondrial fission adaptor proteins are largely species specific. In budding yeast, Fis1 does not directly bind to the dynamin GTPase (Dmn1) at the fission sites, but needs a cytosolic adaptor protein Mdv1 (or its paralogs Caf4) (8). In Apicomplexan parasites, Fis1 is the only identifiable mitochondrial fission protein via bioinformatics (9); BLAST searches of the malaria parasite genome database (www.PlasmoDB.org) did not find any homologs of other mitochondrial fission adaptor proteins in human or yeast (Mff, MiD49, MiD51 or Mdv1/Caf4). Overall, Fis1 seems to the only evolutionarily conserved mitochondrial fission adaptor protein throughout eukaryotic kingdoms.

Most Fis1 homologs are small single transmembrane proteins with ∼ 150 amino acids: the C-terminus contains a transmembrane domain for anchoring onto the MOM and a short post transmembrane tail, whereas the N-terminus has two tetratricopeptide repeats (TPR1/TPR2) that facilitate protein-protein interactions (10). The hypothetical Fis1 in *P. falciparum* (Pf3D7_1325600, 141 aa) shares 30% sequence identity to human Fis1. A modeled 3D structure of PfFis1 with the software I-Tasser (11) is also superimposable with the human Fis1 crystal structure (PDB, 1PC2) (data not shown). In asexual blood stages, PfFis1 is transcribed at all stages with a peak transcription in the late trophozoite and early schizont stages (12), consistent with its expected role in mitochondrial fission. However, it has remained unknown if Fis1 is essential for mitochondrial fission in malaria parasites. The Fis1 homolog in the rodent malaria parasite *Plasmodium berghei* was not included in the large scale gene knockout (KO) study (https://plasmogem.sanger.ac.uk/) (13). The recent PiggyBac mutagenesis survey of *P. falciparum* could not unequivocally assign a phenotype to PfFis1 with statistical confidence due to the short length of the gene (< 500 bp) (14).

In this study, we generated a conditional PfFis1 knockdown (KD) line and a PfFis1 KO line via CRISPR/Cas9 mediated genome editing. In both KD and KO lines of PfFis1, parasites grew normally without noticeable defects, indicating that PfFis1is not essential for mitochondrial fission. We also discovered the important role of the short C-terminal tail of PfFis1 in its correct subcellular localization.

## PfFis1 is localized to the parasite mitochondrion but a conditional knockdown of PfFis1 does not cause defects in the parasite

In order to detect the localization of PfFis1 in *P. falciparum*, we episomally expressed tagged PfFis1 fusion proteins with small epitopes in D10 wildtype (WT) parasites. In one transgenic line, we tagged PfFis1with 3HA at the N-terminus (Fig. 1A) whereas in another transgenic line, PfFis1 was tagged with 3Myc at the C-terminus (Fig. 1B). In both parasites lines, episomal expression of PfFis1 was driven by the promoter of a mitochondrial gene, the 5’-UTR of the annotated mitochondrial ribosomal protein L2 (PfmtRPL2, PF3D7_1132700) (15). Similar to PfFis1, PfmtRPL2 also exhibits a peak transcription at the late trophozoite and schizont stages (www.PlasmoDB.org). Expression of tagged PfFis1 at the predicted molecular weight was confirmed by western blotting (Fig. 1A and 1B). To verify the subcellular localization of PfFis1, we performed immunofluorescence assays (IFA) in both transgenic parasites. Tagged with 3HA at the N-terminus, PfFis1 was localized as expectedly to the mitochondrion; however, PfFis1 was diffused to the cytosol when tagged with 3Myc at the C-terminus (Fig. 1A and 1B). It has been shown recently that truncation of the entire transmembrane domain of Fis1in another apicomplexan parasite, *Toxoplasma gondii*, resulted in mislocalization of the protein in cytoplasm (16), indicating that Fis1 is anchored to MOM via the transmembrane domain. Here, we kept the PfFis1 transmembrane domain intact but merely added 3Myc C-terminally, also resulting in mislocalization of PfFis1. Our data highlights the critical role of the short C-terminal tail (KSFKYF) in protein trafficking and localization of PfFis1 onto the MOM.

**Figure 1.**
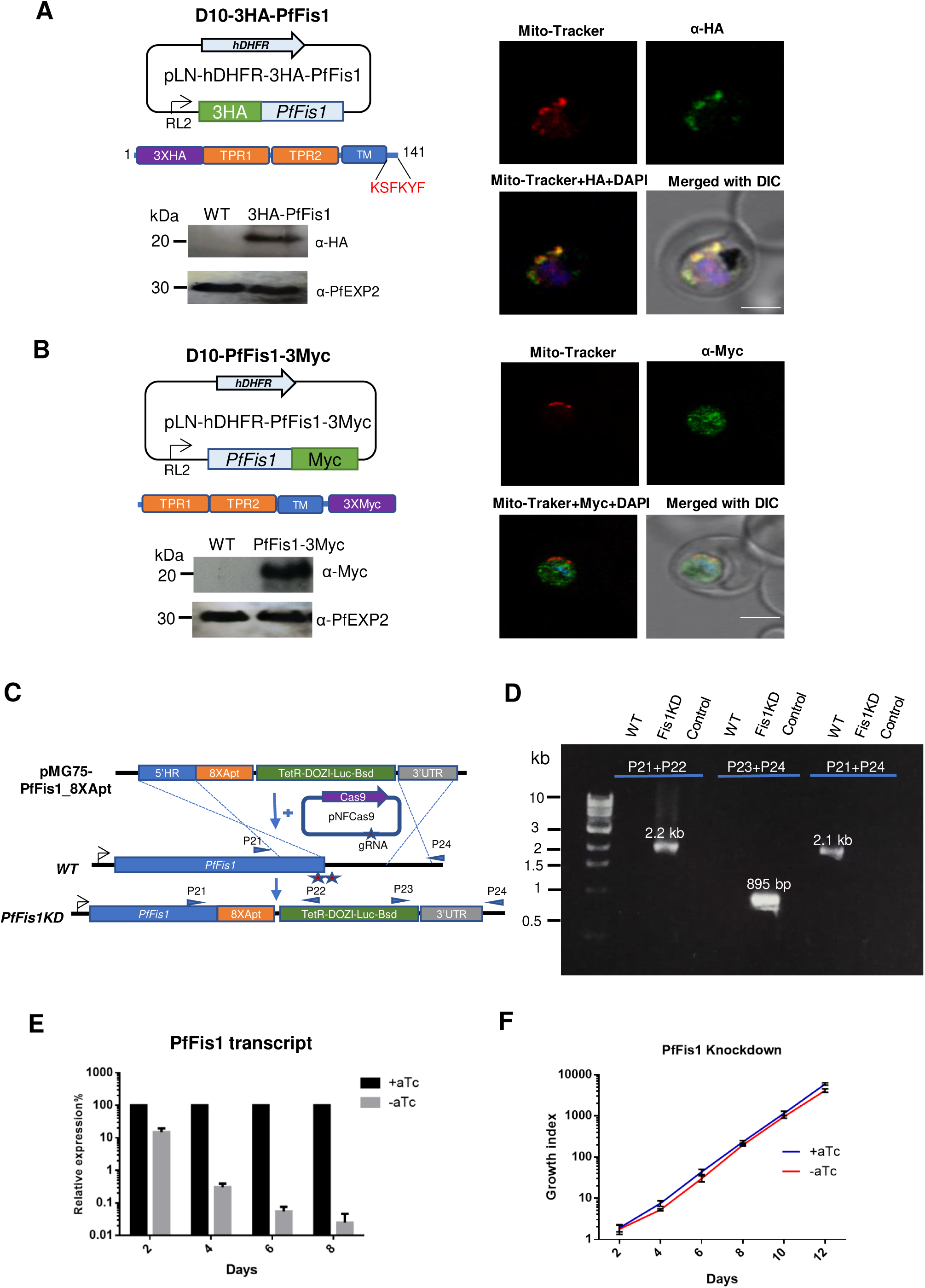
PfFis1 localization with N-or C-terminal tagging and PfFis1 KD without tags. (A) The pLN based construct used for ectopic expression of N-terminal 3HA tagged PfFis1. The C-terminal sequence of PfFis1 (KSFKYF) is highlighted in red. Representative IFA images showing localization of 3HA-PfFis1. Parasites were probed with mouse anti-HA (green), Mito-Tracker (red), DAPI (blue) and merged with DIC, scale bar equals 2µm. Western blot showing the expression of 3HA-PfFis1. Anti-HA and anti-PfEXP2 (loading control) antibodies were used. (B) The pLN based construct used for ectopic expression of C-terminal 3Myc tagged PfFis1. Representative IFA images showing localization of PfFis1-3Myc. Parasites were probed with mouse anti-Myc (green), Mito-Tracker (red), DAPI (blue) and merged with DIC, scale bar equals 2µm. Western blot showing the expression of PfFis1-3Myc. Anti-Myc and anti-PfEXP2 (loading control) antibodies were used. (C) CRISPR/Cas9 based system used to integrate the TetR-DOZI-aptamer system at the 3’ end of the endogenous PfFis1 gene. Positions of primers used in (D) are highlighted. (D) Diagnostic PCR showing the correct genotype of the PfFis1 KD line after genome editing. The PCR product of primers P21+P22 shows the 5’ integration; the PCR product of primers P23+P24 shows the 3’ integration; the PCR product of primers P21+P24 shows the WT genotype. Primers P21+P24 failed to amplify any bands from the KD parasites due to the large size of the fragment to be amplified (>11 kb). Control PCRs contained no template DNA. (E) Analysis of PfFis1 transcripts in the KD parasites via qRT-PCR. At each timepoint, the PfFis1 transcription level in the aTc minus culture was compared to that of the aTc plus culture (the latter was normalized to 100%). Seryl-tRNA synthetase was used as an internal control. Error bars indicate standard deviations from triplicate samples; this experiment has been repeated two times. (F) Effect of PfFis1 KD on the growth of *P. falciparum*. To quantify the growth, parasites were enriched by Percoll, equal number of PfFis1KD parasites was grown in the presence and absence of aTc in the medium (+aTc and - aTc) for 12 days (6 IDCs). Parasitemia was counted on every alternate day and the parasitemia was multiplied by the parasite dilution factor to produce a cumulative measure of growth. All data points are mean±s.d. of three independent experiments.

To determine the role of PfFis1 in parasite survival, we first utilized the CRISPR/Cas9 mediated (17) TetR-DOZI-aptamer system (18) to conditionally knock down the endogenous expression of PfFis1. The conditional KD system is beneficial to evaluate the essentiality of malarial genes as the parasite maintains a haploid genome in the blood stages where knockouts of essential genes are unachievable. In addition, since our data revealed the importance of C-terminus of PfFis1 in its correct localization, we modified our conventional KD systems (19, 20) to reduce the expression of PfFis1, but no tags were added to the C-terminus of PfFis1 (Fig. 1C). We co-transfected WT *P. falciparum* (D10 strain) with the template plasmid, pMG75noP-Fis1-8apt, and two gRNA constructs that were expected to guide Cas9 cleavage near the end of PfFis1 locus. The transfected parasites were selected using media containing blastidicin (Bsd) and anhydrotetracycline (aTc), yielding a conditional KD line. The small molecule aTc maintains the expression level of the targeted gene by preventing the negative regulator, TetR-DOZI fusion protein, from binding to the targeted mRNA which has been tagged by RNA aptamers (18). Hence, expression of PfFis1 was expected to be maintained in aTc supplemented media but abrogated upon aTc removal. The genotype of the PfFis1 KD line was confirmed by diagnostic PCRs using site specific primers (Fig.1D). Since no antibodies were available to detect PfFis1 protein, we verified the KD efficiency by qRT-PCR to quantify PfFis1mRNA transcripts in parasites upon aTc removal for 2, 4, 6 and 8 days (up to 4 IDCs). In comparison to aTc plus controls, the PfFis1 transcript dramatically decreased after aTc removal for 1 IDC (2 days) and continued to decrease to a negligible level over the KD time course (Fig. 1E). With or without aTc, however, the parasites did not show any defects in the growth rate (Fig. 1F), indicating the dispensable role of PfFis1 in parasite survival. Interestingly, our data also revealed that in the absence of aTc, the TetR-DOZI-aptamer system not only prevents protein translation by pulling the mRNA away from ribosomes (18), but also cause a rapid degradation of the target mRNA. To our best knowledge, this is the first report to show that the TetR-DOZI-aptamer system could also interfere with mRNA stability of the target gene.

## PfFis1 is dispensable for mitochondrial fission

Our data thus far has shown that PfFis1 is likely non-essential for the parasite. To rule out the possibility that a low amount of PfFis1 protein in the KD parasite was sufficient to maintain parasite health, we attempted to knock out PfFis1 via CRISPR/Cas9 genome editing. In D10 WT, we co-transfected the KO template plasmid carrying two homologous sequences of PfFis1 with two gRNA constructs that guide Cas9 cleavage in the middle of the PfFis1 genetic locus (Fig. 2A). The transfected parasites were selected by WR99210, a specific inhibitor of the Plasmodium dihydrofolate reductase gene (21) and drug resistant parasites were achieved. The genotype of the transgenic parasite was confirmed by diagnostic PCRs using site specific primers (Fig. 2B). PfFis1 was successfully knocked out, yielding a PfFis1 KO line (PfFis1KO). We tightly synchronized both PfFis1KO and WT lines and observed the growth kinetics over 6 IDCs through parasitemia counting in blood smears of the cultures. PfFis1KO parasites did not exhibit any noticeable growth defects when compared to WT (Fig. 2C), nor did they appear morphologically abnormal (Fig. 2D), suggesting that a complete removal of PfFis1 did not cause problems for parasite growth and replication. To monitor mitochondrial morphologies of PfFis1KO parasites, we stained them with a fluorescent dye Mito-Tracker and performed live cell microscopy. In comparison to WT controls, the mitochondrion of PfFis1KO displayed normal development during the asexual blood stage; importantly, it underwent fission in the late schizont stage to produce fragmented mitochondria to be distributed into daughter cells (Fig. 2E). In addition, we mixed equal numbers of PfFis1KO and WT parasites in one flask, obtained DNA samples every 2 IDCs (4 days) and detected the presence of PfFis1KO parasites by PCR (Fig. 2F). Only PfFis1KO contained the exogenous hDHFR gene (human dihydrofolate reductase gene, the transfection selectable marker) and robustness of the PCR band persisted throughout 16 IDCs (32 days), suggesting that there was no or negligible fitness cost associated with a complete deletion of PfFis1. Collectively, our data suggest that PfFis1 is not essential for mitochondrial fission in the asexual blood stage of *P. falciparum*.

**Figure 2.**
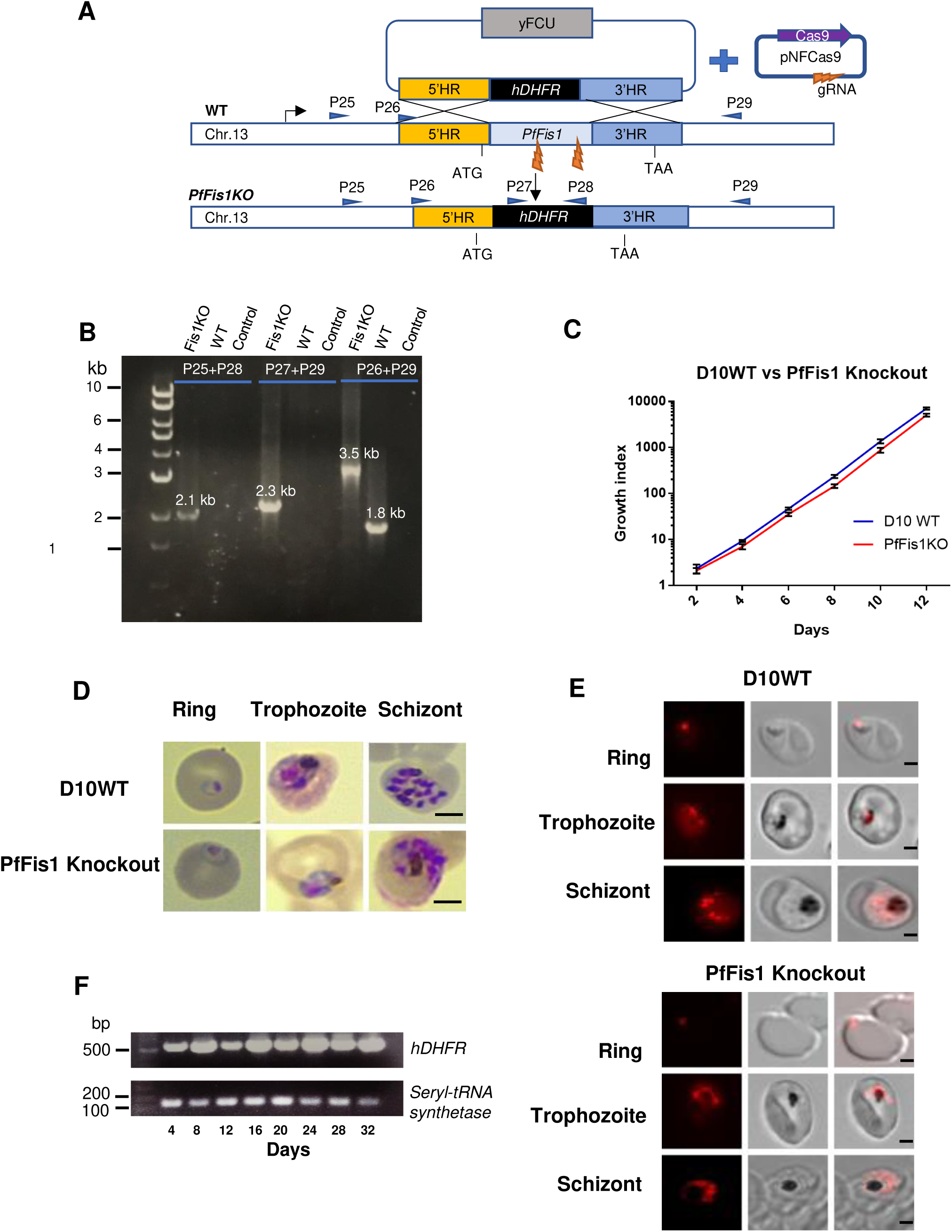
PfFis1 KO does not affect mitochondrial fission. (A) A schematic representation showing the CRISPR/Cas9 mediated replacement of PfFis1 ORF with the hDHFR cassette. Positions of primers used in (B) are highlighted. (B) Diagnostic PCRs showing the correct genotype of the PfFis1 KO line after genome editing. The PCR product of primers P25+P28 shows 5’ integration; the PCR product of primers P27+P29 shows 3’ integration; the PCR product of primers P26+P29 shows the entire locus. The KO band (3.5 kb) is bigger than the WT band (1.8 kb) due to insertion of the hDHFR cassette. (C) Growth curve analysis of PfFis1 KO line compared to D10 WT. Both parasite lines were tightly synchronized and diluted to an initial parasitemia of 1% at ring stages and monitored by analyzing Giemsa-stained blood smears over 12 days. All data points are mean±s.d. of three independent experiments. (D) Morphology of the PfFis1KO parasites analyzed using Giemsa stained blood smears. Scale bars equal 2μm. (E) Live parasite analysis of mitochondrial morphologies. Live PfFis1 KO and D10 WT parasites were treated with Mito-Tracker (10 nM) for 30 min and washed three times. Images were acquired at ring, trophozoite and schizont stages. Scale bars equal 2μm. (F) Growth competition between PfFis1KO and D10 WT parasites. The house-keeping gene seryl-tRNA synthetase was used as an internal control.

### Conclusions

Our data support that PfFis1 relies on its short C-terminal tail for mitochondrial localization, but is not essential for mitochondrial fission or parasite survival in *P. falciparum*. Our Fis1 KO data in malaria parasites is consistent with the proposed non-essentiality of Fis1 by KD approaches carried out in *Toxoplasma gondii* (22). Discovered first in yeast in 2000 (23), Fis1 has been evolutionary conserved in most eukaryotes that contain mitochondria. In yeast, Fis1 is the only known MOM-bound fission adaptor protein and it is essential for mitochondrial fission (24). In human cells, the role of Fis1 in recruiting Drp1 is less critical since several other MOM-bound fission adaptor proteins (Mff, MiD49, MiD51) are capable of playing the same function (6, 25). Interestingly in parasitic apicomplexan parasites that contain significantly smaller genomes, we and others (16, 22) suggest that their mitochondria likely harbor additional MOM-bound fission adaptor proteins to mediate mitochondrial fission. Hence, we propose the mitochondrial fission model in apicomplexan parasites likely resembles that of human cells rather than that of yeast. Further studies are required to identify additional mitochondrial fission adapter proteins in different genus of the Apicomplexa phylum. In particular, the novel and essential mitochondrial fission adaptor protein of apicomplexan parasites could represent potential anti-parasitic drug targets as they are likely absent in the host mitochondria.

## Methods and materials

### Parasite culture and transfection

*Plasmodium falciparum* D10 WT was used for all transfections in this study. Parasite culture and transfection procedures followed our previously published protocols (19, 20).

### Plasmid construction

To tag PfFis1 with 3Myc at the C-terminus, the plasmid pLN-hDHFR-PfFis1*-*3Myc was constructed by amplifying the PfFis1 gene from *P. falciparum* genomic DNA using primers P1+P2. The PCR product was digested with AvrII and BsiWI and inserted into the expression vector pLN-hDHFR-3Myc (19). For N-terminal tagging of PfFis1 with 3HA, a synthetic DNA fragment (AvrII-3HA-BsiWI-PfFis1-AflII) was ordered from Genewiz; the sequence of the DNA fragment is:
tatttttttttgttaatattatacaatataCCTAGGAAAAATGTATCCATACGACGTTCCTGACTATGCCGGATACCCATACGACGTGCCTGATTACGCCGGTTCTTACCCTTACGACGTTCCTGACTACGCCGCACAACGTACGATGGATAGTCCAGAATTACTTAAAATAGAACTTCAAAGATTAAAGAATGATTATGAAAATGAACTATCAGTAGATCACGTAATGCCCAAGACTCAATTTGATTACGCTTGTTTGTTAATATGTTCTTCAGATTTGAAGAACATAAAGTTCGCTTCTTCATTGTTGCATGAATTGTTGTTCATAAATTATAATCGTATAGATTGTTTATATCAGCTAGCTATAGCACATATAAAATTAAGAGATTATAAAAAAGCTAAGAATTATTTAAATGCCTTATTAAAAATCGATGCAAGAAATAGTAATGCTTTAGCTTTAAAGAGTTTACTTTTTGATTTAATATCATCTGATGGTTTAATTGGTGCTTTGTTAGTTGCACTCACAGCTTGTGGTTTATATTTATCTTTTAAATCTTTCAAGTATTTTTAActtaaggtcgagttatataatatatttatgtactcg.

The first 30 and the last 36 base pairs of this synthetic DNA fragment (small letters) are homologous to the ends of the pLN-hDHFR-3HA construct after restriction digestion with AvrII and AflII sites (19). The homologous sequences between the synthetic DNA and the digested vector allowed them to be joined by Infusion reaction (NEBuilder, New England Biolabs).

For both CRISPR/Cas9 mediated KD and KO studies, guide RNAs (gRNAs) were identified from the gene sequence using the Eukaryotic Pathogen CRISPR guide RNA design Tool (http://grna.ctegd.uga.edu/). They were individually cloned into the NFCas9-yDHOD(-) plasmid which contains a full length Cas9 gene from *Streptococcus pyogenes* without any tags and a gRNA expression cassette (20). The gRNA sequences are listed in P3-P6. Diagnostic PCRs were used to identify positive clones from Infusion reactions. These primers are listed in P7-P10. The reverse primer P11 was used as previous described (20). All gRNA constructs were sequenced by Genewiz using primer P12.

For constructing the template plasmid for KD studies without tags, the 5’HR (5’ homologous region) of PfFis1 was amplified from genomic DNA by primers P13+P14. The 3’UTR (3’ homologous region) of PfFis1 was amplified from genomic DNA by primers P15+P16. The 3’UTR was cloned into the pMG75noP-8apt-3HA vector (20) by AflII and BspeI, whereas the 5’HR fragment was subsequently cloned by BspeI and ApaI, yielding the plasmid pMG75noP-Fis1-8apt for KD. This plasmid was linearized with EcoRV before transfection. For constructing the template plasmid for KO studies, the 5’HR was amplified from genomic DNA by primers P17+P18. The 3’HR was amplified from genomic DNA by primers P19+P20. The two HR fragments were sequentially cloned into pCC1, yielding the plasmid pCC1-5’3’Fis1 for KO. This plasmid was linearized with HincII before transfection. Primers of P21-P24 were used to check the genotype of the KD line; primers of P25-P29 were used to check the KO genotype.

### Parasite growth kinetics, IFA, Western blotting, live Mito-Tracker staining

These procedures were followed according to our published protocols (19, 20).

### Nucleic acid extraction, PCR and qRT-PCR

Genomic DNA from late-stage parasites was isolated with DNeasy Blood and Tissue kit (Qiagen). During the KD time course (days 2, 4, 6 and 8), total RNA from parasites of each condition (aTc plus vs aTc minus) was isolated from saponin lysed parasite pellets followed by treatment with Trizol (Thermo) and purification with RNeasy kit (Qiagen). After treated with DNase I (New England Biolabs), 2 µg RNA of each condition was primed with random hexamers and converted to cDNA using SuperScript III reverse transcriptase (Thermo). qRT-PCR was carried out in triplicate with SYBR Green Real-Time PCR Master Mixes (Thermo) in the Real-time PCR instrument (Applied Biosystems). Primers used for amplification of PfFis1 are listed as P30-P31. A previously reported house-keeping gene, seryl-tRNA synthetase, was used as the internal control (primers P32-P33) (26). Data was analyzed using 2^-ΔΔCt^ method as previously described (27). For regular PCR, a reaction volume of 25 µL was used and extension temperature of the Taq polymerase was kept at 62°C.

### Primers used in this study

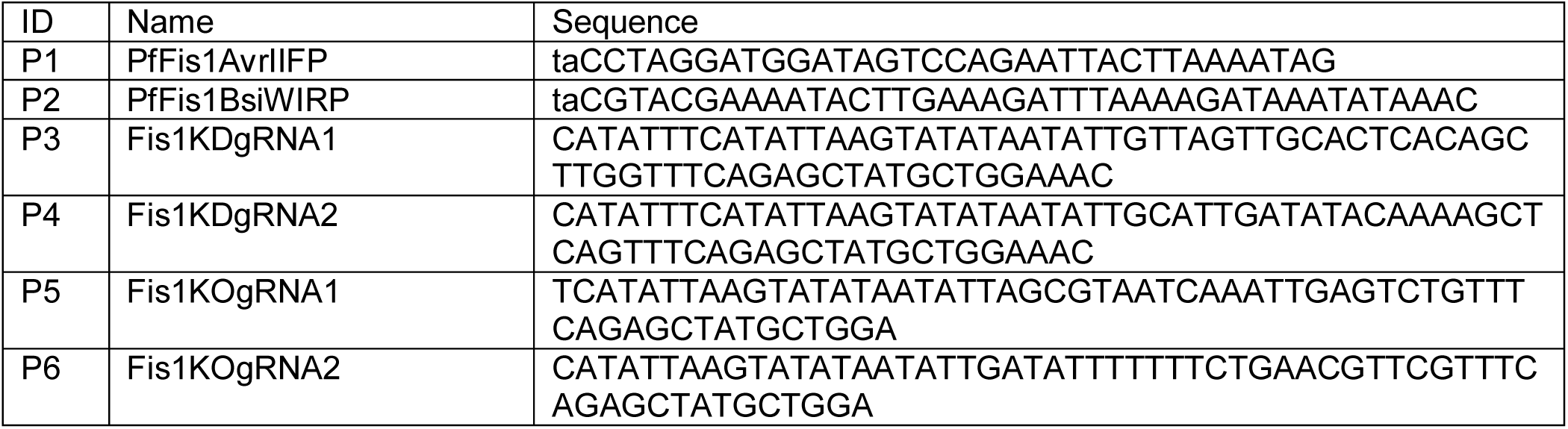
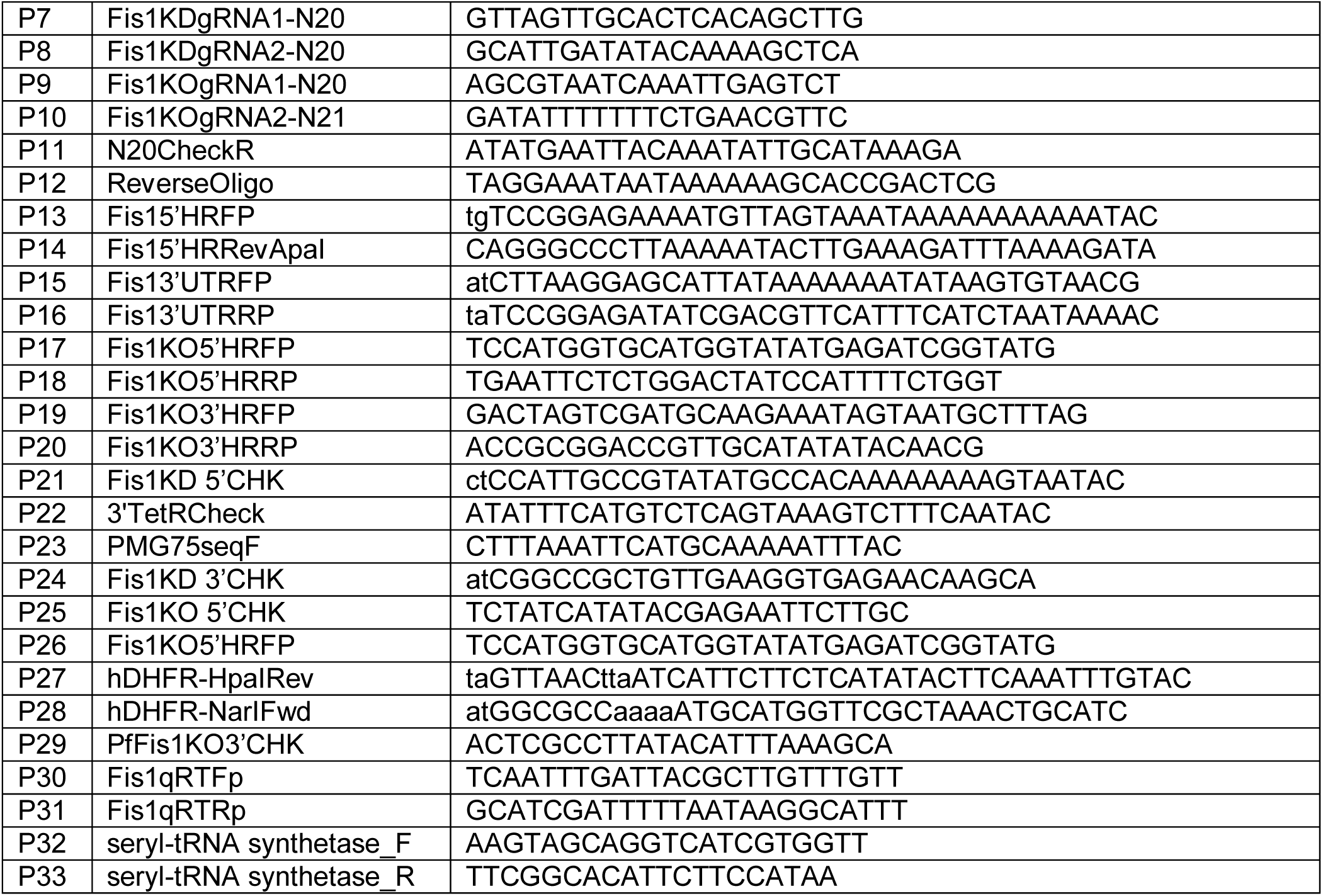

## Conflict of interest

The authors declare no conflicts of interest with the contents of this article.

## Acknowledgements

We thank Dr. Josh Beck (Iowa State University, United States), Dr. Jacquin Niles (MIT, United States), and Dr. James Burns (Drexel University, United States) for sharing plasmid vectors and reagents. We thank the members of the Center for Molecular Parasitology in the Department of Microbiology and Immunology at Drexel University College of Medicine for constructive discussion of this study.

## Funding

This work was supported by an NIH career transition award (K22, K22AI127702) to Dr. Hangjun Ke.

